# Development of an Ethylenediaminetetraacetic Acid-Enhanced Deep Proteomic Profiling Method for Dried Blood Spots and its Application in Mouse Disease Models

**DOI:** 10.64898/2026.07.13.738354

**Authors:** Daisuke Nakajima, Toshio Kanno, Yusei Okuda, Hiromasa Mitsui, Ryo Konno, Naho Ueyama, Yusuke Endo, Osamu Ohara, Yusuke Kawashima

## Abstract

Dried blood spots (DBS) are well-established microsamples used in clinical testing and newborn screening. However, their use in deep proteomics is hindered by highly abundant blood proteins and inefficient protein recovery from filter paper matrices. Notably, the non-targeted analysis of non-specifically DBS-absorbed proteins (NANDA) workflow partially overcomes the impact of abundant blood proteins and has enabled the identification of over 5,000 proteins from DBS. Nonetheless, residual abundant proteins, including hemoglobin and fibrinogen, constrain deep proteomic analysis. Therefore, this study aimed to evaluate the effects of the metal chelator ethylenediaminetetraacetic acid (EDTA) on the depth of DBS proteomic analysis. An optimized EDTA-enhanced NANDA protocol that incorporated a 100 mM EDTA wash step was compatible with standard DBS collection procedures and required no modification of current clinical workflows, reducing residual fibrinogen-chain signals and facilitating its potential use in clinical and translational settings. When combined with the Orbitrap Astral data-independent acquisition mass spectrometry, this approach enabled the single-shot identification of more than 7,000 proteins from DBS. To the best of our knowledge, this represents the deepest proteome coverage reported for DBS to date. The workflow also supports high-throughput and highly reproducible analyses. Additionally, application of this workflow to mouse disease models revealed disease-specific systemic immune signatures from minimal blood volumes. Collectively, these results establish EDTA-enhanced NANDA as a practical and scalable workflow that overcomes longstanding limitations of DBS proteomics, thereby enabling deep, high-throughput, minimally invasive proteomic profiling across diverse biological and experimental contexts.

---

Dried blood spots (DBS) are widely used in clinical testing, epidemiological studies, and newborn screening (NBS) because they require a small volume of blood and offer sample stability and ease of transportation.^1–5^ Owing to their minimally invasive and straightforward collection procedure, DBS are useful biospecimens when blood volume is limited or sample transportation is difficult. Such settings include large-scale cohort studies and remote healthcare settings.^6^

Recent advances in mass spectrometry-based proteomics have enabled comprehensive profiling of protein expression in the blood, thereby enabling potential applications in disease diagnosis and biomarker discovery. However, technical challenges remain for deep proteomic profiling of DBS. Specifically, blood contains highly abundant proteins, such as hemoglobin and albumin, which may hinder the detection of less abundant proteins. Furthermore, inefficient extraction of proteins adsorbed onto filter paper matrices may limit the proteome coverage of DBS.^7^

As an alternative microsampling technology, volumetric absorptive microsampling (VAMS) has recently attracted attention.^8,9^ In particular, VAMS enables the collection of a defined blood volume using an absorptive device, thereby reducing variability in the sample volume. Accordingly, proteomic analyses using VAMS have identified more than 2,000 proteins in blood samples. However, VAMS requires specialized devices, which increase costs and may limit its widespread adoption.

In contrast, owing to its low cost and simple sampling procedure, DBS sampling remains a globally established sampling method within the existing NBS infrastructure and remains an essential microsampling approach. To improve DBS proteomics, we previously developed a sodium carbonate precipitation method to reduce the impact of highly abundant blood proteins. This enabled the identification of approximately 2,000 proteins from DBS.^10–12^ More recently, we developed the non-targeted analysis of non-specifically DBS-absorbed proteins (NANDA) workflow, which exploits differences in protein retention on filter paper during aqueous washing.^13^

In this workflow, DBS filter paper is dispersed into a pulp-like suspension, followed by sequential washing. Proteins remaining associated with the filter paper pulp are then directly subjected to enzymatic digestion, and the resulting peptides are recovered for LC–MS/MS analysis. Importantly, the NANDA workflow does not selectively retain proteins according to their abundance. Instead, it exploits differences in protein solubility and retention on the filter paper matrix during aqueous washing. In general, water-soluble proteins, such as hemoglobin and albumin, tend to be more readily removed, whereas relatively poorly soluble proteins, including membrane-associated and cytoskeletal proteins, tend to remain associated with the filter paper. Cellulose, the major component of DBS filter paper, is generally hydrophilic owing to its abundant hydroxyl groups, although cellulose surfaces can also exhibit amphiphilic properties.^14^ Protein adsorption at the cellulose–liquid interface has been demonstrated in studies of blood droplets deposited on paper.^15^ Previous studies have shown that protein adsorption onto cellulose depends on the physicochemical properties of individual proteins and the surrounding solution conditions, including pH and ionic strength. Multiple noncovalent interactions, including hydrophobic interactions, van der Waals forces, and hydration-related effects, may contribute to this adsorption behavior.^16^ In addition to these surface interactions, the porous fibrous structure of filter paper may contribute to the physical retention of poorly soluble proteins and cellular components. Therefore, protein retention during NANDA washing likely reflects a combination of protein solubility, adsorption to the cellulose matrix, and physical retention, although the relative contributions of these mechanisms remain unclear. Although this separation is not absolute, the removal of readily extractable abundant proteins reduces their contribution to the resulting peptide mixture, thereby facilitating the detection of proteins with lower signal intensities. This approach enabled the identification of more than 5,000 proteins from DBS.^13^ To facilitate automated processing, we further developed an iron powder-assisted NANDA workflow (auto-NANDA), in which iron powder serves as a magnetic carrier for collecting and transferring the filter paper pulp between washing solutions without centrifugation. Using a robotic magnetic-handling system, this workflow enables the simultaneous processing of 96 DBS samples within approximately 1 h, excluding subsequent enzymatic digestion and peptide cleanup.^13^ The basic principle and automated sample preparation procedure of NANDA are illustrated in Figure 1A.

**Figure 1.**
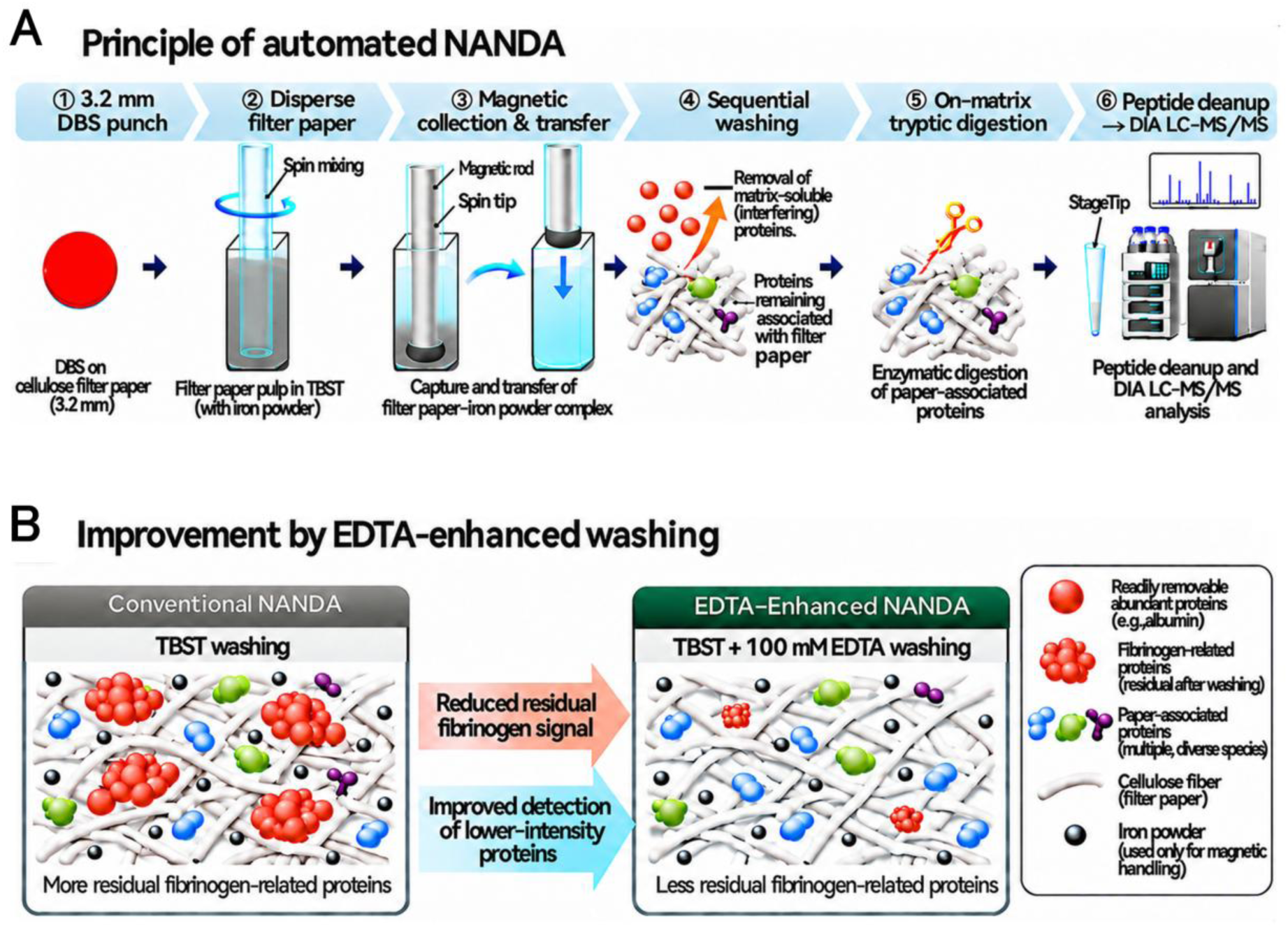
Principles of automated NANDA and EDTA-enhanced NANDA for DBS proteome analysis. (A) Overview of the automated NANDA workflow. A 3.2-mm DBS punch is mechanically dispersed into a pulp-like suspension in TBST containing iron powder. A magnetic rod inserted into a spin tip enables collection and transfer of the filter paper–iron powder complex between plates. Sequential washing removes readily extractable proteins, followed by on-matrix tryptic digestion of proteins remaining associated with the washed filter paper pulp. The resulting peptides are purified using an SDB StageTip and analyzed by DIA LC–MS/MS. Iron powder serves primarily as a magnetic carrier for handling the filter paper pulp. (B) Schematic comparison of conventional NANDA and EDTA-enhanced NANDA. Incorporation of 100 mM EDTA into the TBST washing buffer reduces the residual fibrinogen-related signal and improves the detection of lower-intensity proteins. The schematic illustrates the observed analytical effects without establishing the molecular mechanism underlying the EDTA effect. Abbreviations: DBS, dried blood spot; EDTA, ethylenediaminetetraacetic acid; TBST, Tris-buffered saline with Tween 20; SDB, styrene– divinylbenzene; DIA, data-independent acquisition; LC–MS/MS, liquid chromatography–tandem mass spectrometry.

However, further improvements in proteome depth are required to fully realize the analytical potential of DBS proteomics. In particular, a more effective suppression of highly abundant blood proteins is required to enhance the detection of low-abundance species. Accordingly, in this study, we evaluated ethylenediaminetetraacetic acid (EDTA), a widely used anticoagulant and metal-ion chelator, both during DBS preparation and as an additive to the NANDA washing buffer to determine its effects on residual abundant protein signals and proteome coverage. The application during the washing step yielded a greater number of identified proteins, was fully compatible with standard DBS collection procedures, and required no modification of existing clinical workflows. The differences between conventional NANDA and EDTA-enhanced NANDA are schematically illustrated in Figure 1B. Furthermore, the use of EDTA could have implications for research in animal models. Specifically, considering that DBS are prepared from small volumes of blood, repeated blood sampling can be performed in mice without sacrificing the animals. This will enable longitudinal monitoring of disease progression or treatment responses. In addition, although hemolysis is often difficult to avoid during blood collection from small animals, DBS are prepared from whole blood absorbed onto filter paper. Therefore, these samples are less affected by hemolysis. Thus, deep proteomic profiling using DBS has the potential to provide a minimally invasive and comprehensive approach for investigating systemic molecular changes in small-animal disease models.

Therefore, this study aimed to establish an EDTA-enhanced NANDA workflow. The improved workflow was further applied to mouse DBS to assess its applicability to small-animal disease models.

## EXPERIMENTAL PROCEDURES

### Preparation of Human DBS

Human DBS were obtained from consenting healthy volunteers using a finger prick on blood sampling paper (Advantec Toyo Kaisha, Ltd., Tokyo, Japan; cat. no. PKU-S). The samples were dried overnight at room temperature (20–25 °C). DBS from anticoagulated blood were prepared using whole blood collected in tubes containing sodium heparin (Terumo Corporation, Tokyo, Japan; cat. no. VO-H050K) and sodium citrate (Terumo Corporation; cat. no. VP-J053K), or EDTA-2K (Terumo Corporation; cat. no. VP-DK050K). Thereafter, 60 µL of whole blood was dispensed onto each circle (diameter: 1 cm) of sampling paper without centrifugation using a micropipette and dried overnight. The study involving human participants was approved by the Institutional Review Board of the Kazusa DNA Research Institute (approval no. 2020-06). Written informed consent was obtained from all participants, and the study was conducted in accordance with the Declaration of Helsinki.

### Preparation of EDTA-Impregnated Filter Paper

One edge of the filter paper was immersed in 5, 50, 100, 200, or 500 mM EDTA solution, which permeated the paper via capillary action. The filter paper was subsequently air-dried thoroughly.

### Preparation of Mouse DBS

Peripheral blood was collected from the mice via the lateral tail vein using a minimally invasive procedure. Mice (C57BL/6 background) were gently restrained using a mouse restrainer (Braintree Scientific, Braintree, MA, USA), and the tail was warmed under a heat lamp (∼37 °C) for 1–2 min to dilate the veins. The skin was disinfected with 70% ethanol, and a 25-gauge needle (Terumo Corporation) was used to puncture the lateral tail vein at approximately one-third of the base of the tail. Approximately 20 µL of blood was collected via capillary action into EDTA-coated capillary glass tubes (Paul Marienfeld GmbH & Co. KG, Lauda-Königshofen, Germany; cat. no. 2909000). Subsequently, the blood was blotted onto blood sampling paper and dried overnight. After blood collection, hemostasis was achieved in mice by applying a sterile gauze with gentle pressure for 1–2 min. The mice were monitored until they fully recovered and returned to their cages.

### Mouse Disease Model

The following two immune-mediated disease models with distinct characteristics were selected: experimental autoimmune encephalomyelitis (EAE)^17,18^ and a vitamin D3 analogue MC903 (MCE# HY-10001)-induced atopic dermatitis (AD) model.^19^ EAE reflects Th1/Th17-driven central nervous system inflammation, whereas MC903-induced AD represents Th2-dominant skin inflammation. Both models exhibit systemic immune changes detectable in blood, which renders them suitable for evaluating DBS-based proteomic analysis. To establish the EAE model, mice were immunized subcutaneously at two dorsal sites on day 0 with 100 µg MOG35-55 peptide emulsified in complete Freund’s adjuvant. This was followed by intraperitoneal injection of 100 ng pertussis toxin on days 0 and 1 according to the manufacturer’s protocol (Hooke Laboratories, Lawrence, MA, USA). The clinical signs of EAE were assessed as follows: 0, no disease; 0.5, partially limped tail; 1, limped tail; 2, limped tail and partial hind leg paralysis; 3, complete hind leg paralysis; 4, tetraparesis; and 5, moribund. Mice were euthanized when their scores reached ≥4. For the AD model, MC903 was dissolved in ethanol (EtOH) and topically applied on mouse ears (2 nmol in 20 µL per ear) daily for 9 days. The same volume of EtOH was used as the vehicle control. The animal experiments were performed according to protocols approved by the Institutional Animal Care and Use Committee of the Kazusa DNA Research Institute (Registration No. 30-1-002). Animal care was performed according to the guidelines of the Kazusa DNA Research Institute.

### Protein Extraction from DBS for Proteome Analysis

Protein preparation from DBS was performed using our previously established automated NANDA (auto-NANDA) workflow.^13^ Briefly, the workflow consists of mechanical disruption of DBS filter paper into a pulp-like suspension in the presence of iron powder, magnetic collection and transfer of the filter paper–iron powder complex, sequential washing to remove readily extractable proteins, and subsequent enzymatic digestion of proteins remaining associated with the filter paper. Iron powder primarily serves as a magnetic carrier for handling the filter paper pulp rather than as a protein-selective affinity reagent. The EDTA-enhanced NANDA workflow developed in this study follows the same overall procedure but incorporates EDTA into the washing step (Figure 1A, B).

For automated processing, 96-well plates compatible with a Maelstrom 9610 instrument (TANBead, Taoyuan City, Taiwan) were prepared. Thereafter, 500 µL of Tris-buffered saline with Tween 20 (TBST; 25 mM Tris, 137 mM NaCl, 2.68 mM KCl, 0.05% Tween 20, pH 7.4; Nacalai Tesque, Kyoto, Japan), 5 mg of freshly prepared iron powder suspension (particle size: 3–5 µm; Kojundo Chemical Lab. Co., Ltd., Japan; 5 mg/25 µL in 50% glycerol), and one disk (diameter: 3.2 mm) punched from the DBS were added into the first plate. The second plate was filled with 500 µL of TBST, whereas the third and fourth plates each received 500 µL of 50 mM Tris-HCl (pH 8.0). The fifth plate was filled with 200 µL of 50 mM Tris-HCl (pH 8.0), 10 mM CaCl_2_, and 0.012% laurylmaltose neopentylglycol (LMNG; Anatrace Products, LLC, Maumee, OH, USA).^20^ In the EDTA-enhanced workflow, the TBST solutions in both the first and second plates contained 100 mM EDTA (final concentration); the third, fourth, and fifth plates contained no added EDTA. The plate configuration and processing sequence are shown in Figure 2. For the EDTA concentration-dependence experiment, the EDTA concentration in the TBST solutions was varied as indicated in Figure 6A–C.

**Figure 2.**
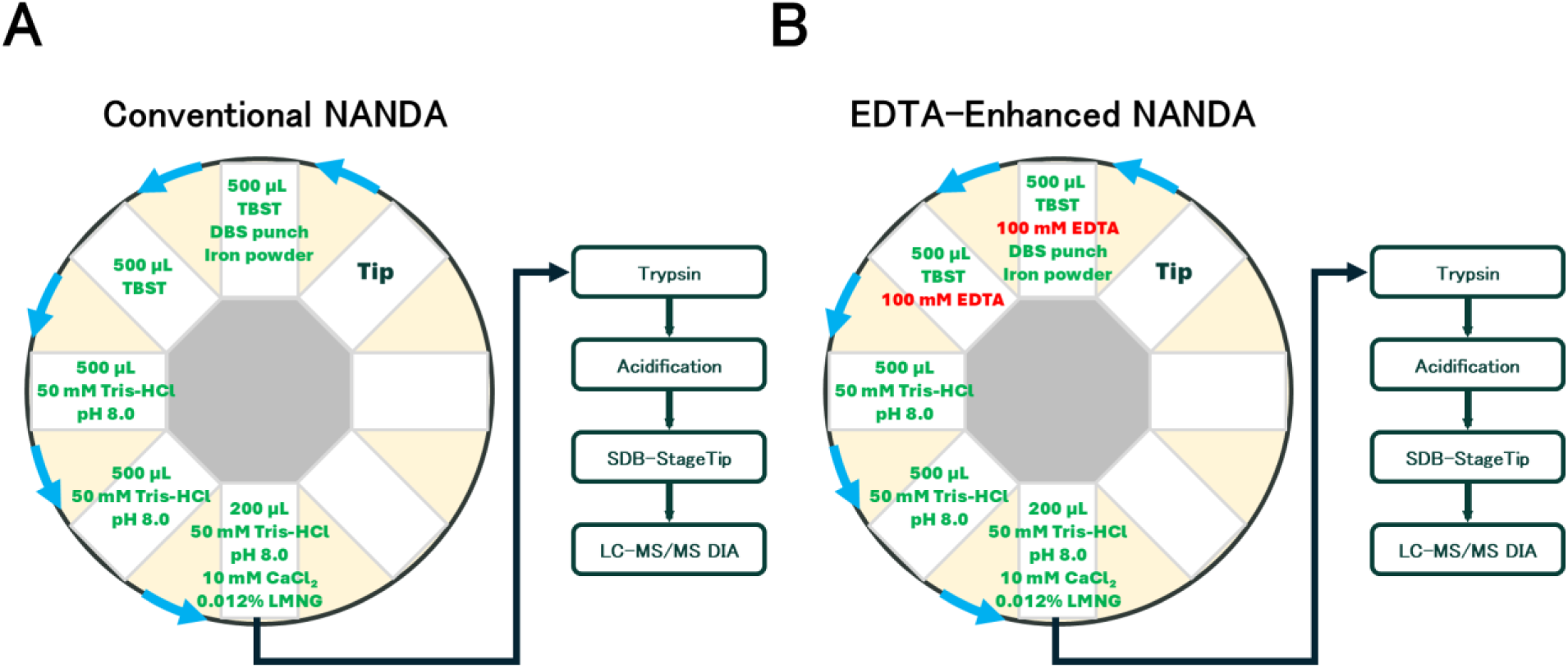
Comparison of conventional and EDTA-enhanced automated NANDA workflows for DBS proteome analysis. (A) Conventional NANDA. A 3.2-mm DBS punch is mechanically dispersed in 500 µL of TBST containing 5 mg of iron powder in the first plate. The filter paper–iron powder complex is magnetically collected and sequentially transferred through a second plate containing 500 µL of TBST and third and fourth plates each containing 500 µL of 50 mM Tris–HCl (pH 8.0). (B) EDTA-enhanced NANDA. The same procedure is used, with 100 mM EDTA added to both the initial dispersion solution and the subsequent TBST wash in the first and second plates. In both workflows, the washed complex is transferred into a fifth plate containing 200 µL of 50 mM Tris–HCl (pH 8.0), 10 mM CaCl₂, and 0.012% laurylmaltose neopentylglycol (LMNG). Proteins remaining associated with the washed filter paper pulp are digested with 1 µg of trypsin for 14 h at 37 °C, as described in the Experimental Procedures. Following removal of the filter paper residue, the supernatant is acidified with TFA, and peptides are purified using an SDB StageTip before DIA LC–MS/MS analysis. Arrows indicate the sequence of magnetic transfers and downstream processing. Abbreviations: DBS, dried blood spot; EDTA, ethylenediaminetetraacetic acid; TBST, Tris-buffered saline with Tween 20; TFA, trifluoroacetic acid; SDB, styrene–divinylbenzene; DIA, data-independent acquisition; LC–MS/MS, liquid chromatography–tandem mass spectrometry.

Subsequently, the plates were mounted on the Maelstrom 9610, and protein extraction from DBS was initiated. In this system, a spin tip is used to agitate the sample in each well, while a magnetic rod inserted into the spin tip enables magnetic collection and transfer of the iron powder-containing complex between wells. A spin tip was inserted into the first plate, and the DBS was disrupted via agitation in TBST for 30 min at 3,500 rpm with inversion every 10 s, thereby forming a pulp-like DBS–iron powder complex. A magnetic rod was then inserted into the spin tip, and the complex was captured and collected on the tip for 60 s. This was followed by a transfer to the second plate containing TBST. The magnetic rod was then withdrawn, allowing the collected complex to be resuspended in the washing buffer. The complex was washed via agitation for 10 min at 3,500 rpm with inversion every 10 s. Subsequently, the complex was transferred to the third plate containing 50 mM Tris-HCl (pH 8.0) and washed under the same agitation conditions for 10 min. This washing step was repeated in the fourth plate containing 50 mM Tris-HCl (pH 8.0), with agitation for 2 min. Finally, the complex was transferred to the fifth plate containing 50 mM Tris-HCl (pH 8.0), 10 mM CaCl_2_, and 0.012% LMNG and suspended via agitation for 2 min at 3,500 rpm with inversion every 10 s.

### Protein Digestion

For on-matrix digestion of proteins remaining associated with the washed filter paper pulp, 1 µg of Trypsin Platinum (cat. no. VA9000; Promega, Madison, WI, USA) was gently mixed with the sample and incubated for 14 h at 37 °C. After removing the filter paper residue from the digested sample using Maelstrom 8-Autostage/Maelstrom 9610, the isolated supernatant was acidified with 50 µL of 5% trifluoroacetic acid (TFA). Thereafter, the sample was desalted using a styrene– divinylbenzene polymer stop- and-go extraction tip (GL Sciences, Tokyo, Japan), which was washed with 25 µL of 80% acetonitrile (ACN) in 0.1% TFA. This was followed by equilibration with 50 µL of 3% ACN in 0.1% TFA. Subsequently, the sample was loaded onto the tip, washed with 80 µL of 3% ACN in 0.1% TFA, and eluted with 50 µL of 36% ACN in 0.1% TFA. A 5 µL aliquot of the eluate was used for peptide quantification using the Pierce Quantitative Fluorometric Peptide Assay Kit (Cat. No. 23290; Thermo Fisher Scientific, Waltham, MA, USA). Briefly, 5 µL of the peptide sample was mixed with 35 µL of Fluorometric Peptide Assay Buffer and 10 µL of Fluorometric Peptide Assay Reagent. After incubation at room temperature for 5 min, fluorescence was measured at excitation and emission wavelengths of 390 and 475 nm, respectively. Peptide concentrations were determined using an eight-point standard curve (0–500 µg/mL) prepared from the Peptide Digest Assay Standard supplied with the kit. The standards and samples were analyzed using identical reagent volumes and reaction conditions, maintaining the manufacturer’s recommended mixing ratio of 1:7:2 (sample:buffer:reagent). The remaining 45 µL of the eluate was dried in a centrifugal evaporator (miVac Duo Concentrator; Genevac, Ipswich, UK). Based on the measured peptide concentration, the dried peptides were reconstituted at 100 ng/µL in 0.1% TFA containing 0.02% decyl maltose neopentyl glycol (DMNG; Anatrace Products, LLC, Maumee, OH, USA).^20^ The reconstituted samples were then transferred to liquid chromatography (LC) vials (Thermo Fisher Scientific, Waltham, MA, USA). The assay uses fluorescent labeling of peptide N-terminal amino groups. No study-specific estimates of assay precision or accuracy are reported. The injection amount was calculated from the measured concentration and injection volume and therefore represents a nominal amount subject to peptide-assay and sample-handling variability.

### LC-Tandem Mass Spectrometry with Data-Independent Acquisition

#### Orbitrap Exploris 480 mass spectrometer analysis (12 samples/day)

A 2 µL aliquot of the peptide solution (100 ng/µL), corresponding to a nominal peptide amount of 200 ng, was injected onto a 75 µm × 30 cm nanoLC column (ReproSil-Pur C18, particle size: 1.7 µm, 100 Å; CoAnn Technologies, Richland, WA, USA). The column was maintained at 50 °C, and peptides were separated using a 90-min gradient comprising 0 min of 6% B, 76 min of 34% B, 83 min of 70% B, and 90 min of 70% B, at a flow rate of 150 nL/min (solvent A: 0.1% formic acid in water; solvent B: 0.1% formic acid in 80% acetonitrile).

Chromatographic separation was performed using an UltiMate 3000 RSLC nanosystem (Thermo Fisher Scientific). Eluted peptides were analyzed using the Orbitrap Exploris 480 mass spectrometer (Thermo Fisher Scientific) equipped with an InSpIon system (AMR, Tokyo, Japan).^21^ For data-independent acquisition (DIA),^22–24^ mass spectrometry (MS1) spectra were acquired over a 495–705 m/z range at a resolution of 60,000, with an automatic gain control (AGC) target of 300% and a maximum injection time set to “Auto.” MS2 spectra were acquired over a 200–1,800 m/z range at a resolution of 60,000, with an AGC target of 3,000%, a maximum injection time set to “Auto,” and stepped normalized collision energies of 22, 26, and 30%. The optimized window was set at 4 Th, and the arrangement was configured using Xcalibur 4.3 (Thermo Fisher Scientific).

#### Orbitrap Astral mass spectrometer analysis (16 samples/day)

A 2 µL aliquot of the peptide solution (100 ng/µL), corresponding to a nominal peptide amount of 200 ng, was injected onto a 75 µm × 30 cm nanoLC column (ReproSil-Pur C18, particle size: 1.5 µm, 100 Å; CoAnn Technologies) at 60 °C. Separation was conducted under the following conditions: an 84.5-min gradient comprising 1% B at a flow rate of 600 nL/min in 0–0.5 min, 1– 8% B at a flow rate of 600–200 nL/min in 0.5–4.5 min, 8–23% B at a flow rate of 200 nL/min in 4.5–60 min, 23–40% B at a flow rate of 200 nL/min in 60–78.5 min, 40–98% B at a flow rate of 200 nL/min in 78.5–79.5 min, 98% B at a flow rate of 200 nL/min in 79.5–80.5 min, 98% B at a flow rate of 200–600 nL/min in 80.5–81.5 min, 98% B at a flow rate of 600–750 nL/min in 81.5– 82.5 min, and 98% B at a flow rate of 750 nL/min in 82.5–84.5 min.

Chromatographic separation was performed on a Vanquish Neo UHPLC system (Thermo Fisher Scientific). Subsequently, the peptides eluted from the column were analyzed using the Orbitrap Astral (Thermo Fisher Scientific) equipped with an InSpIon system. MS1 spectra were collected in the range of m/z 380–980 at a 240,000 resolution using the Orbitrap analyzer to set an AGC target of 500% and a maximum injection time of 5 ms. MS2 spectra were collected at m/z 250– 2,000 using the Astral analyzer to set an AGC target of 600%, a maximum injection time of 4 ms, and a normalized collision energy of 25%. The isolation width of MS2 was set to 1.6 Th.

#### Orbitrap Astral mass spectrometer analysis (50 samples/day)

A 2 µL aliquot of the peptide solution (100 ng/µL), corresponding to a nominal peptide amount of 200 ng, was injected onto a 75 µm × 30 cm nanoLC column (ReproSil-Pur C18, particle size: 1.5 µm, 100 Å; CoAnn Technologies) at 60 °C. Separation was conducted under the following conditions: a 23-min gradient comprising 1% B at a flow rate of 700 nL/min in 0–0.5 min, 1–10% B at a flow rate of 700–400 nL/min in 0.5–3.5 min, 10–28% B at a flow rate of 400 nL/min in 3.5– 15.5 min, 28–42% B at a flow rate of 400 nL/min in 15.5–19.5 min, 42–98% B at a flow rate of 400 nL/min in 19.5–20.5 min, 98% B at a flow rate of 400 nL/min in 20.5–21.5 min, 98% B at a flow rate of 400–800 nL/min in 21.5–22.5 min, and 98% B at a flow rate of 800 nL/min in 22.5– 23 min.

Chromatographic separation was performed on a Vanquish Neo UHPLC system (Thermo Fisher Scientific). Subsequently, the peptides eluted from the column were analyzed using the Orbitrap Astral (Thermo Fisher Scientific) equipped with an InSpIon. MS1 spectra were collected in the range of m/z 380–980 at a 240,000 resolution using the Orbitrap analyzer to set an AGC target of 300% and a maximum injection time of 5 ms. MS2 spectra were collected at m/z 200–2,000 using the Astral analyzer to set an AGC target of 300%, a maximum injection time of 3 ms, and a normalized collision energy of 25%. The isolation width for MS2 was set to 2 Th.

### Data Analysis

For DBS protein analysis, DIA-MS data were queried against species-specific *in silico* spectral libraries using DIA-NN v.2.2.0 (https://github.com/vdemichev/DiaNN).^25^ Initially, a spectral library was generated from the human protein sequence UniProt database (proteome ID UP000005640, 20,644 entries, downloaded on March 28, 2025) or the UniProt mouse proteome database (proteome ID UP000000589, 21,755 entries, downloaded on April 9, 2025) using DIA-NN. The parameters for generating the spectral library were as follows: digestion enzyme, trypsin; missed cleavage, 1; peptide length range, 7–45; precursor charge range, 2–4; and fragment ion m/z range, 200–1,800 for Orbitrap Exploris 480 and 200–2,000 for Orbitrap Astral. The precursor mass range varied according to the DIA method (500–700 m/z and 380–980 m/z). Additionally, “FASTA digest for library-free search/library generation,” “deep learning-based spectra, RTs and IM prediction,” and “n-term M excision” were enabled. For the DIA-NN search, the following parameters were applied: mass accuracy and MS1 accuracy were set according to the instrument used (10 and 10 ppm, respectively, for Orbitrap Exploris 480 data; 7 and 4 ppm, respectively, for Orbitrap Astral data), scoring in peptideforms, proteotypicity based on genes, machine learning in NNs (cross-validated), quantification was conducted using QuantUMS (high precision), and cross-run normalization was set to “RT-dependent.” Furthermore, “Unrelated runs” and “Protein inference” were enabled. Match-between-runs (MBR) was disabled for all analyses except those used for Figures 7E and S2B, for which MBR was enabled. Mouse protein identification counts (Figure 8A and Table S2-1) were obtained by analyzing individual runs independently. The mouse dataset used for Figure 8B–H and Table S2-2 was generated by jointly analyzing all 12 runs with cross-run normalization and without MBR. The threshold for protein identification was set to ≤1% for the precursor and protein false discovery rates.

Human proteins are labeled using gene symbols approved by the HUGO Gene Nomenclature Committee (HGNC). Mouse proteins are labeled using the corresponding mouse gene symbols, displayed in uppercase in the main text and Figure 8G,H. The protein quantification values were aggregated over the quantification values of the unique peptides, as calculated using the DIA-NN. Log2 transformation of protein intensity was performed, and a filtering step was conducted to ensure that at least one group for each protein contained ≥70% valid values. Missing values were imputed using random numbers drawn from the normal distribution (width, 0.3; downshift, 1.8) using Perseus v1.6.15.0,^26^ and Pearson’s correlation analysis was performed. The criterion for identifying altered proteins included an absolute fold change of ≥2 and a Welch’s *t*-test *p*-value of <0.05 between the two groups. Disease-associated protein coverage was assessed by matching identified proteins to the Online Mendelian Inheritance in Man (OMIM) database,^27^ the Recommended Uniform Screening Panel (RUSP),^28^ and the recessive-gene panel of Building Evidence and Collaboration for GenOmics in Nationwide Newborn Screening (BEACONS; https://www.beaconsnbs.org/). Gene Ontology (GO) enrichment analyses for molecular function (MF), biological process (BP), and cellular component (CC) of the 773 human proteins detected exclusively with the 100 mM EDTA wash were performed using DAVID (https://david.ncifcrf.gov/tools.jsp), with the default *Homo sapiens* genome-wide gene set provided by DAVID as the background. The top 10 terms in each category, ranked by *p*-value, are presented in Figure S1A–C, respectively. Graphs were generated using GraphPad Prism 9.5.1 (GraphPad Software, San Diego, CA, USA) and Microsoft Excel (Microsoft Corporation, Redmond, WA, USA).

### Experimental Design and Statistical Rationale

All quantitative data are presented as mean ± standard deviation, unless otherwise specified. The number of replicates and the unit of comparison are specified in the corresponding figure legends. Human DBS comparisons used three or four measurements per condition, as indicated; the mouse disease-model comparisons included four mice per group. The reproducibility analysis in Figure 7E comprised LC–MS/MS measurements of 16 independently prepared DBS samples processed using the EDTA-enhanced NANDA workflow. Missing values were imputed using random numbers drawn from a normal distribution (width = 0.3, downshift = 1.8) using Perseus. Pearson’s correlation analysis was subsequently performed to assess relationships between samples. Statistical comparisons between two groups were conducted using a two-tailed unpaired Welch’s *t*-test. Proteins were considered significantly altered if they exhibited an absolute fold change of ≥2 and a Welch’s *t*-test *p*-value of <0.05. Statistical significance was defined as follows: *p* < 0.05 (*), *p* < 0.01 (**), *p* < 0.001 (***), and not significant (ns). All statistical analyses were conducted using Perseus or GraphPad Prism.

## RESULTS AND DISCUSSION

### Impact of Blood Coagulation on Deep Proteome Analysis of DBS

During DBS preparation, blood was applied to filter paper and allowed to dry at room temperature. Coagulation reactions may occur before complete drying. Blood coagulation is a Ca²⁺-dependent process in which fibrinogen is converted to fibrin, forming an insoluble fibrous network. Such a network could contribute to protein retention within the filter paper matrix; however, fibrin formation and its effects on protein resolubilization were not directly measured in this study.

All human DBS method-development experiments shown in Figures 3–6 were analyzed by DIA LC–MS/MS using the Orbitrap Exploris 480 at 12 samples per day (SPD). Similarly, in NANDA, when high-abundance proteins were removed and the remaining proteins were analyzed, the highly intense proteins among the identified set were predominantly hemoglobin and fibrinogen (Figure 3). The efficient removal of hemoglobin and fibrinogen enhances the detection of low-abundance proteins; therefore, DBS were prepared using anticoagulated blood to investigate whether their removal from the NANDA system could be improved. Accordingly, DBS were generated from whole blood collected in anticoagulant-containing tubes (heparin, citrate, and EDTA) and subjected to proteomic analysis using NANDA. DBS prepared from blood collected with any of the three anticoagulants showed reduced residual fibrinogen-chain signals after NANDA washing (Figure 4A), consistent with inhibition of fibrin formation before drying. This reduction was particularly pronounced with citrate and EDTA.

**Figure 3.**
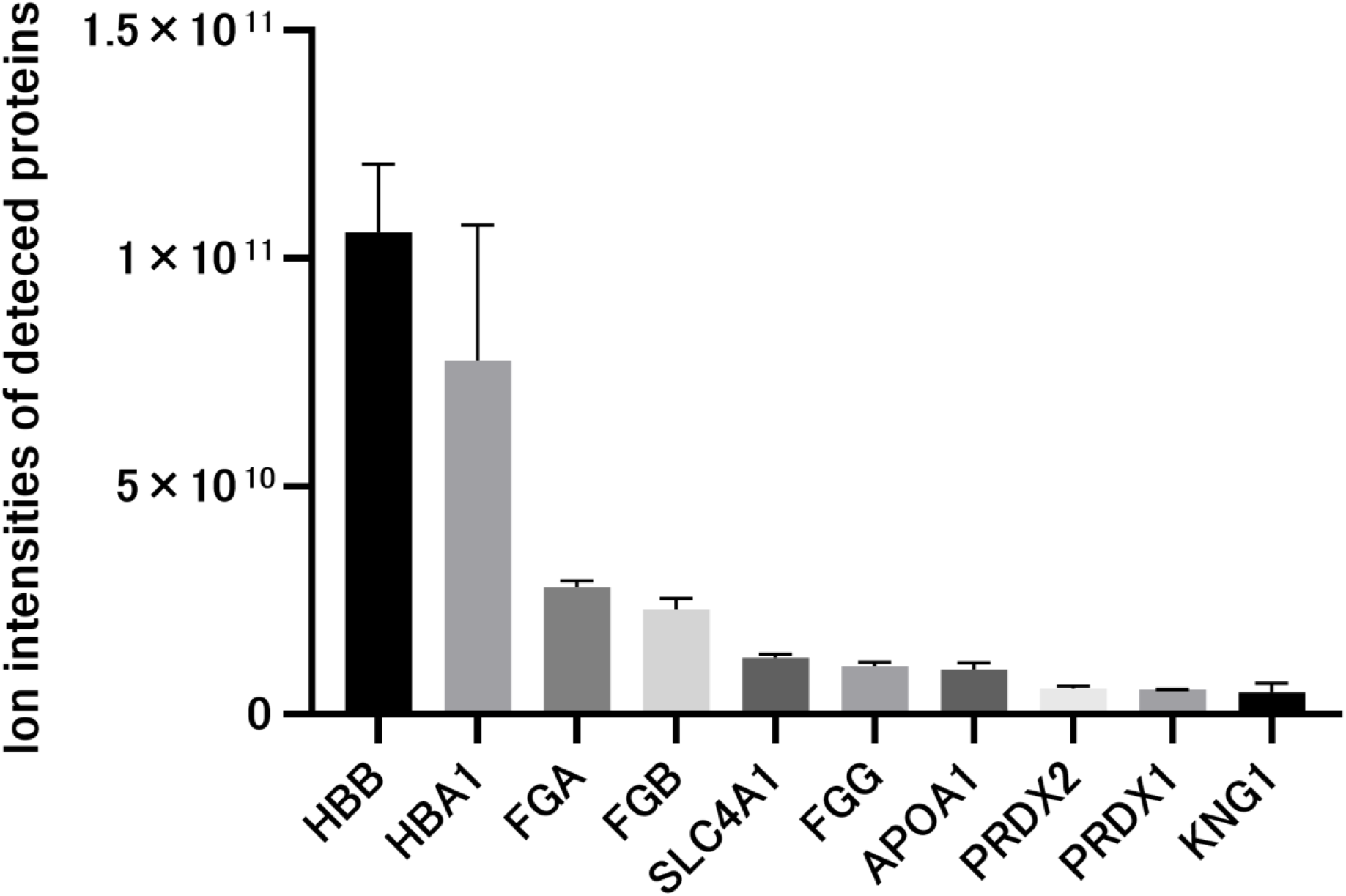
Ion intensities of the 10 most abundant proteins detected in DBS using automated NANDA (auto-NANDA). Data are shown as the mean ± SD (*n* = 4). Proteins are labeled using HUGO Gene Nomenclature Committee (HGNC)-approved gene symbols. DIA LC–MS/MS analysis was performed using an Orbitrap Exploris 480 at 12 SPD. Abbreviations: DBS, dried blood spot; NANDA, non-targeted analysis of non-specifically DBS-absorbed proteins; HBB, hemoglobin subunit beta; HBA1, hemoglobin subunit alpha 1; FGA, fibrinogen alpha chain; FGB, fibrinogen beta chain; SLC4A1, solute carrier family 4 member 1; FGG, fibrinogen gamma chain; APOA1, apolipoprotein A1; PRDX2, peroxiredoxin-2; PRDX1, peroxiredoxin-1; KNG1, kininogen-1; SD, standard deviation; SPD, samples per day.

**Figure 4.**
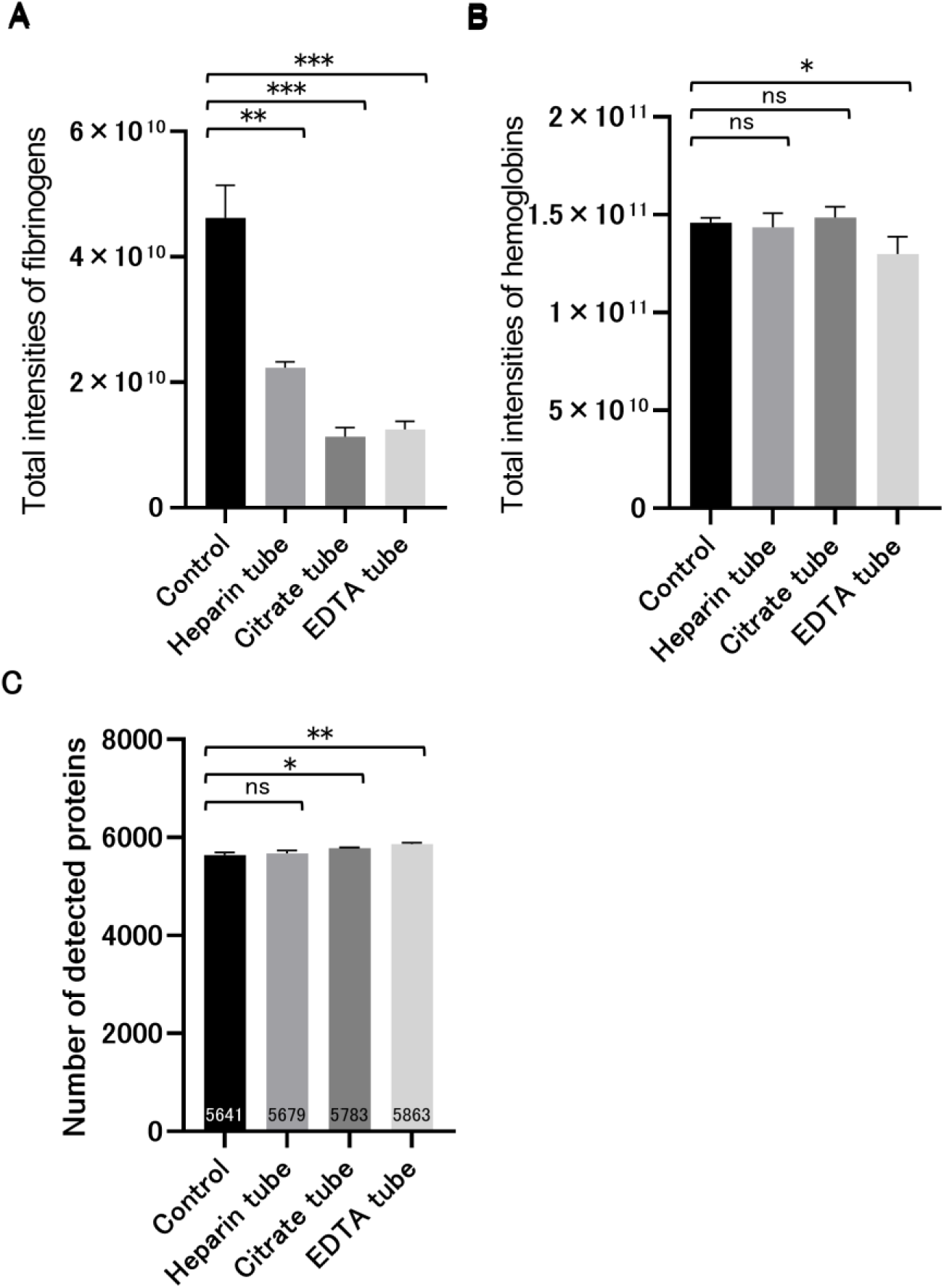
Comparison of protein detection in DBS prepared using different blood collection procedures. (A) Summed ion intensities of fibrinogen chains (FGA, FGB, and FGG). (B) Summed ion intensities of hemoglobin subunits (HBA1, HBB, HBD, HBG1, HBG2, HBM, HBQ1, and HBZ). (C) Number of detected proteins, shown as the mean ± SD of protein counts for individual samples (*n* = 4 per group). Control DBS samples were prepared by directly spotting blood onto filter paper. The other DBS samples were prepared from blood collected in tubes containing heparin, citrate, or EDTA. DIA LC–MS/MS analysis was performed using an Orbitrap Exploris 480 at 12 SPD. Statistical significance was assessed using a two-tailed, unpaired Welch’s *t*-test: \**p* < 0.05; \*\**p* < 0.01; \*\*\**p* < 0.001; ns, not significant. Abbreviations: DBS, dried blood spot; EDTA, ethylenediaminetetraacetic acid; SD, standard deviation; SPD, samples per day.

Moreover, a slight reduction in hemoglobin was observed with EDTA treatment (Figure 4B). In contrast, when blood collection tubes containing heparin or citrate were used, no marked improvement in hemoglobin removal efficiency was observed, despite their anticoagulant properties. These results suggest that inhibition of blood coagulation alone may not fully explain the lower residual hemoglobin signal observed with EDTA. EDTA chelates divalent metal ions, including Ca²⁺, Mg²⁺, and Zn²⁺, and its presence during blood collection and drying may influence interactions among blood components or their retention within the filter paper matrix. However, citrate also chelates calcium ions, and the present study did not directly assess changes in metal-dependent protein interactions, hemoglobin solubility, or adsorption to cellulose. Therefore, the precise mechanism responsible for the modest reduction in residual hemoglobin remains unclear.

In total, 5,641 proteins were detected in the conventional DBS prepared by directly spotting blood onto filter paper, whereas 5,679, 5,783, and 5,863 proteins were identified under heparin, citrate, and EDTA conditions, respectively (Figure 4C). The EDTA condition showed the greatest improvement, with a 3.9% increase compared with that of the control. This increase was associated with lower residual fibrinogen-chain signals and a modest reduction in hemoglobin signal; the individual contributions of these changes to proteome coverage were not determined. Although the benefits of using EDTA-containing blood collection tubes were demonstrated, changing DBS blood collection protocols across hospitals for proteome analysis is challenging. Therefore, this study sought to suppress blood coagulation without altering the blood collection procedure by preparing an EDTA-impregnated filter paper in which EDTA was preloaded onto the paper used for blood spotting.

The EDTA-impregnated filter paper was prepared by soaking the paper in 5–500 mM EDTA, followed by drying. Whole blood was spotted onto paper to generate DBS, which were then analyzed using NANDA. Fibrinogen- and hemoglobin-chain signals decreased with increasing EDTA concentration, consistent with an effect of EDTA impregnation on the residual abundant-protein signals (Figure 5A, B). With respect to the number of identified proteins, 5,479 proteins were detected in DBS prepared using normal filter paper. In contrast, the number of identified proteins increased with the EDTA-impregnated filter paper, with 5,781, 5,794, 5,862, 5,914, and 6,179 proteins detected using 5, 50, 100, 200, and 500 mM EDTA, respectively (Figure 5C). These results demonstrate the effectiveness of the EDTA-impregnated filter paper, which is easily prepared at a low cost. Hence, EDTA-impregnated filter paper is feasible for distribution to individual hospitals.

**Figure 5.**
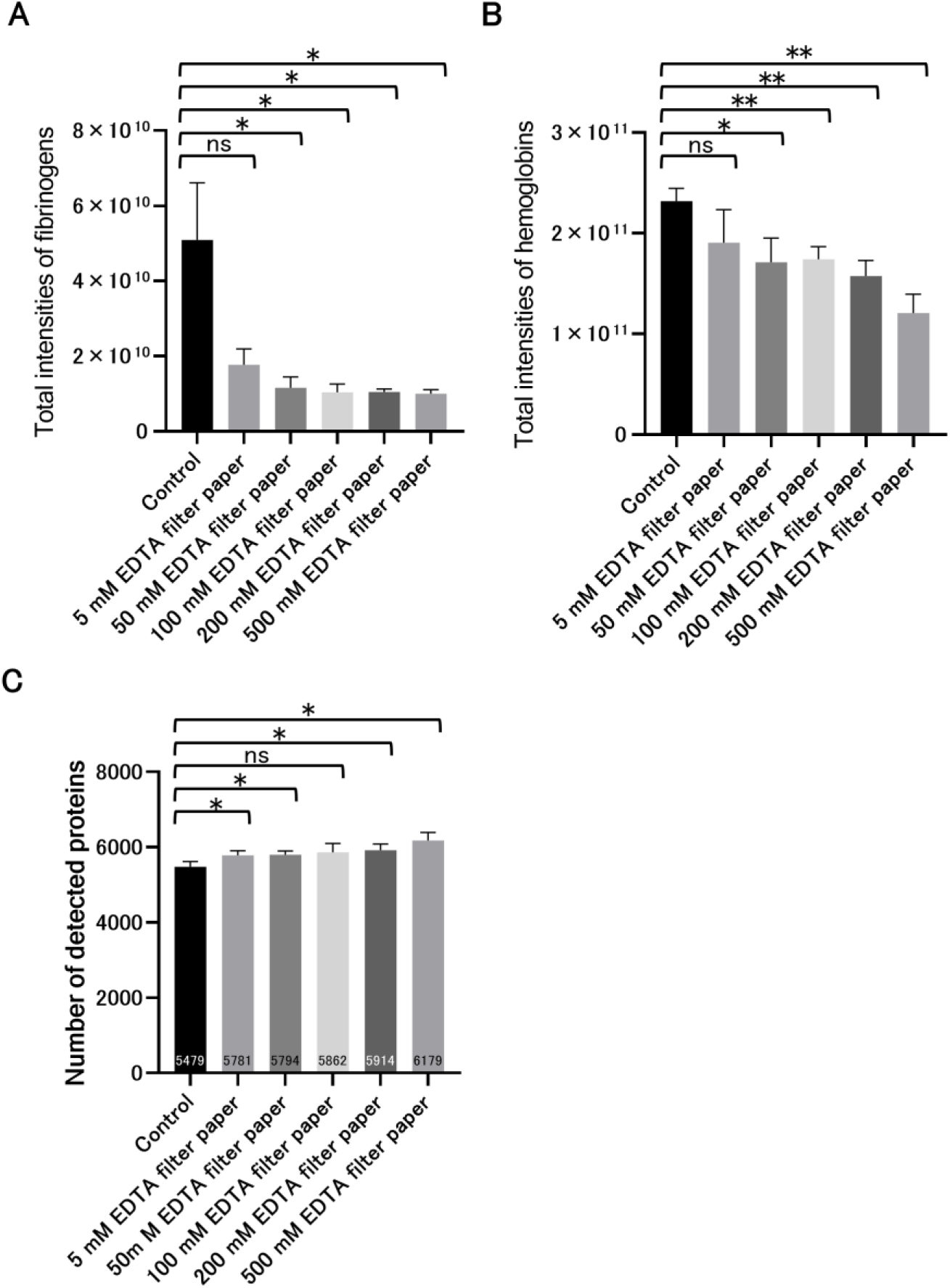
Effect of EDTA-impregnated filter paper on DBS protein detection. (A) Summed ion intensities of fibrinogen chains (FGA, FGB, and FGG). (B) Summed ion intensities of hemoglobin subunits (HBA1, HBB, HBD, HBG1, HBG2, HBM, HBQ1, and HBZ). (C) Number of detected proteins, shown as the mean ± SD of protein counts for individual samples (*n* = 3 per group). DBS samples were prepared using plain filter paper (control) or filter paper impregnated with EDTA solutions at concentrations of 5, 50, 100, 200, or 500 mM. DIA LC–MS/MS analysis was performed using an Orbitrap Exploris 480 at 12 SPD. Statistical significance was assessed using a two-tailed, unpaired Welch’s *t*-test: \**p* < 0.05; \*\**p* < 0.01; ns, not significant. Abbreviations: DBS, dried blood spot; EDTA, ethylenediaminetetraacetic acid; SD, standard deviation; SPD, samples per day.

### Effect of EDTA on Conventional DBS

In contrast to the anticoagulant experiments described above, in which EDTA was present before DBS formation, we investigated whether EDTA could also affect protein retention when introduced only during the washing of conventionally prepared DBS. Specifically, EDTA was added to the washing buffer in the NANDA method, and the residual amounts of hemoglobin and fibrinogen as well as the number of identified proteins were evaluated. Fibrinogen levels decreased in an EDTA concentration-dependent manner (Figure 6A). In contrast, the residual amount of hemoglobin reached a minimum at 50 mM and subsequently increased in a concentration-dependent manner (Figure 6B). The mean numbers of identified proteins were 5,373, 5,713, 5,839, 6,094, 6,056, and 6,120 for control, 25, 50, 100, 250, and 450 mM EDTA, respectively. Protein identification increased up to 100 mM and remained at a similar level at higher concentrations (Figure 6C).

**Figure 6.**
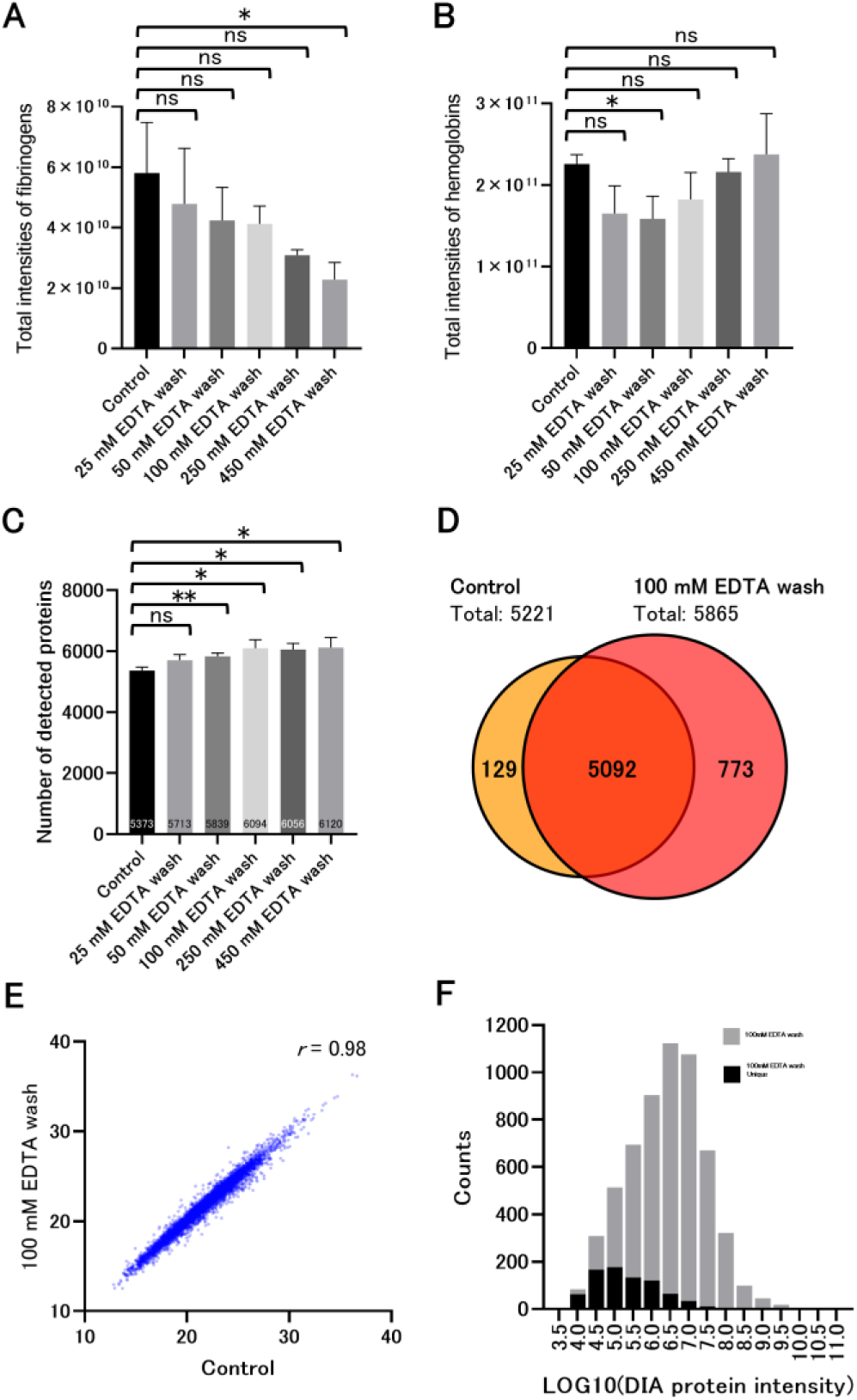
Effect of EDTA supplementation of the wash buffer on DBS protein detection using auto-NANDA. (A) Summed ion intensities of fibrinogen chains (FGA, FGB, and FGG). (B) Summed ion intensities of hemoglobin subunits (HBA1, HBB, HBD, HBG1, HBG2, HBM, HBQ1, and HBZ). (C) Number of detected proteins, shown as the mean ± SD of protein counts for individual samples (*n* = 3 per condition). For panels A–C, DBS samples were washed with TBST alone (control) or TBST containing 25, 50, 100, 250, or 450 mM EDTA during auto-NANDA processing. Statistical significance was assessed against the control using two-tailed, unpaired Welch’s *t*-tests: \**p* < 0.05; \*\**p* < 0.01; ns, not significant. Panels D–F show results from a separate experiment comparing TBST alone (control) with TBST containing 100 mM EDTA (*n* = 4 per condition). (D) Venn diagram comparing the sets of proteins detected in all four replicates within each condition (*n* = 4 per condition). (E) Correlation of protein ion intensities between the control and 100 mM EDTA wash conditions for proteins detected in both conditions. The x- and y-axes represent the mean log₂-transformed protein ion intensities for the respective conditions. (F) Histogram showing the distributions of mean protein ion intensities on a log₁₀ scale for all proteins detected under the 100 mM EDTA wash condition (gray) and proteins uniquely detected under this condition (black). DIA LC–MS/MS analysis was performed using an Orbitrap Exploris 480 at 12 SPD. Abbreviations: DBS, dried blood spot; EDTA, ethylenediaminetetraacetic acid; NANDA, non-targeted analysis of non-specifically DBS-absorbed proteins; auto-NANDA, automated NANDA; TBST, Tris-buffered saline with Tween 20; SD, standard deviation; SPD, samples per day.

The increase in the number of identified proteins was associated with reduced residual signals of fibrinogen chains and, at certain EDTA concentrations, hemoglobin (Figure 6A–C). EDTA is a strong chelator of divalent metal ions, including Ca²⁺, which plays an important role in fibrinogen stability and fibrin assembly.^29,30^ Therefore, EDTA-mediated changes in the metal-ion environment during washing may influence the retention and release of proteins associated with the DBS filter paper matrix. However, calcium chelation does not necessarily disrupt preformed fibrin networks, particularly those stabilized by covalent cross-linking.^31,32^ Furthermore, because the LC–MS/MS measurements were based on peptides derived from fibrinogen chains, they could not distinguish fibrinogen from fibrin. Thus, the observed reduction in fibrinogen-related signals is consistent with improved removal during washing but does not directly demonstrate disruption of the fibrin network. In contrast, hemoglobin exhibited a nonmonotonic response to EDTA concentration, with its lowest residual signal observed at 50 mM. This behavior suggests that additional factors, potentially including changes in the ionic environment and protein–matrix interactions, may influence hemoglobin retention. Further investigation is required to elucidate the precise mechanisms underlying these observations.

A 100 mM EDTA wash was selected as a practical condition that improved proteome coverage and reduced residual fibrinogen-chain signals while preserving compatibility with standard DBS collection. This concentration was not selected to minimize hemoglobin: its lowest residual signal occurred at 50 mM, and higher EDTA concentrations did not provide a clear additional gain in proteome coverage (Figure 6A–C). Figure 6C reports the mean number of proteins identified per sample in a concentration-optimization experiment (*n* = 3 per condition). In a separate experiment (*n* = 4 per condition), proteins detected in all four replicates within each condition were compared. There were 5,092 shared proteins, 129 control-specific proteins, and 773 proteins detected only with the 100 mM EDTA wash, giving condition-specific totals of 5,221 and 5,865 proteins, respectively (Figure 6D). Thus, the counts in Figure 6C and 6D differ because they derive from separate experiments and different counting criteria. GO enrichment analyses of these 773 proteins highlighted MF terms related to ribosomal structure, RNA binding, signaling adaptor activity, and transcriptional regulation (Figure S1A). BP terms included mitochondrial translation, translation, transcription initiation, GPI anchor biosynthesis, and lysosome organization (Figure S1B). CC terms included the nucleoplasm, lysosomal and Golgi membranes, and mitochondrial compartments (Figure S1C). These findings suggest that the EDTA wash extends proteomic coverage to proteins involved in diverse cellular processes and associated with multiple subcellular compartments. The mean log2-transformed intensities of shared proteins were strongly correlated (Pearson *r* = 0.98; Figure 6E). This correlation indicates broad agreement in relative protein abundance but does not exclude protein-specific quantitative differences. Proteins detected only with EDTA were predominantly distributed in the lower-intensity range (Figure 6F), consistent with improved detection of proteins with weaker signals.

To compare the 100 mM EDTA wash with 500 mM EDTA-impregnated filter paper directly, proteins detected in all three replicates within each condition were examined (Figure S2A). Of the 5,706 proteins detected with the EDTA wash and the 5,771 detected with EDTA-impregnated paper, 5,281 were shared; 425 and 490 were specific to the wash and impregnated-paper conditions, respectively. The shared proteins represented 85.2% of the union, demonstrating substantial but incomplete overlap. Pairwise correlations of protein ion intensities across the six samples are shown in Figure S2B. Correlation coefficients were approximately 0.99 within each condition and 0.96–0.97 between conditions, indicating broad quantitative agreement with somewhat lower agreement between the two preparation methods. These results support comparable overall coverage while showing that the detected protein sets are not identical. Although 500 mM EDTA-impregnated paper also reduced residual hemoglobin and fibrinogen signals, its introduction would require a change to the collection material. In contrast, the 100 mM EDTA wash can be applied to conventionally collected DBS, including samples obtained through existing NBS workflows. We therefore selected the wash-based approach for this study and designated it EDTA-enhanced NANDA, without claiming that it is universally superior to EDTA-impregnated paper.

### Proteome Coverage and Reproducibility of EDTA-Enhanced NANDA

The DBS processed using the EDTA-enhanced NANDA workflow were analyzed using a high-performance mass spectrometer (the Orbitrap Astral) to determine the number of proteins that could be identified from DBS in a single-shot analysis. Moreover, the Orbitrap Astral was used to evaluate the reproducibility of the analysis of these proteins using a high-throughput method with potential application to NBS. To maximize protein identification in a single-shot measurement, a method with a throughput of 16 samples per day (SPD) was employed. Compared with the 6,195 proteins identified using the EDTA-enhanced NANDA workflow with the 12 SPD method on the Orbitrap Exploris 480 (Exploris 12 SPD), the 16 SPD method on the Orbitrap Astral (Astral 16 SPD) identified 7,166 proteins (Figure 7A; Table S1). Previous studies have demonstrated progressive improvements in DBS proteome coverage. Chambers et al. identified an average of 253 proteins using untargeted LC–MS/MS,^7^ whereas a sodium carbonate precipitation-based approach enabled the identification of 1,977 proteins.^10^ Subsequently, 2,912 proteins were consistently quantified across healthy newborn DBS samples in a clinical study.^11^ More recently, the automated NANDA workflow enabled the identification of more than 5,000 proteins while supporting a throughput of up to 45 samples per day.^13^ In the present study, the effect of EDTA was directly evaluated under matched LC–MS/MS conditions. On the Orbitrap Exploris 480 at 12 SPD, the number of identified proteins increased from 5,552 with conventional NANDA to 6,195 with EDTA-enhanced NANDA, representing an 11.6% improvement (Figure 7A). Furthermore, combining EDTA-enhanced NANDA with Orbitrap Astral enabled the identification of 7,166 proteins at 16 SPD. Although differences in sample preparation, instrumentation, acquisition time, and protein identification criteria limit direct comparisons across studies, these results demonstrate improved proteome coverage with EDTA washing under matched conditions and highlight the analytical depth achievable with the optimized workflow.

**Figure 7.**
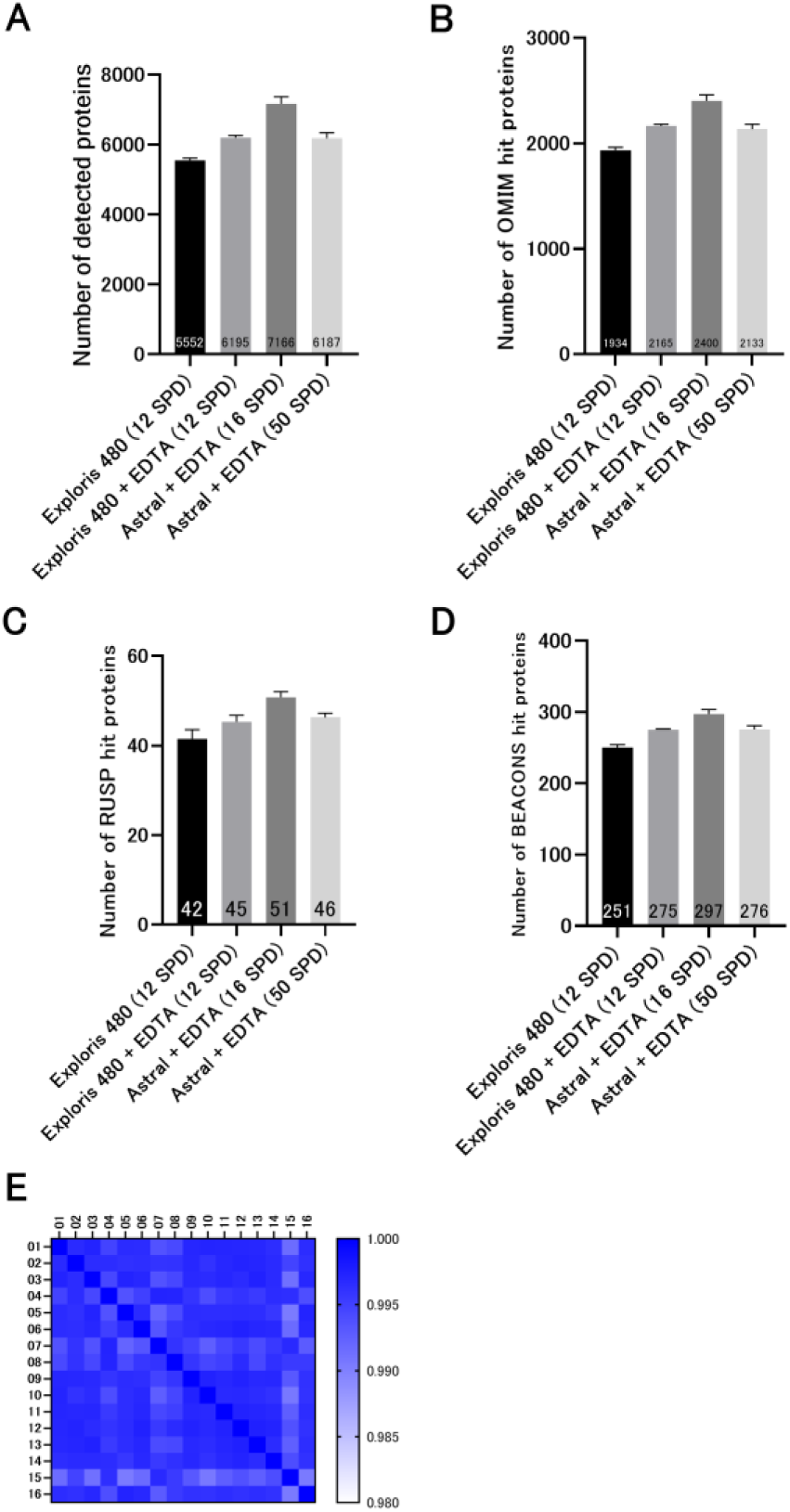
Proteome coverage and reproducibility of DBS analysis using EDTA-enhanced NANDA workflows. (A) Number of proteins detected in DBS using the conventional NANDA workflow with an Orbitrap Exploris 480 (12 SPD) or EDTA-enhanced NANDA workflows with an Orbitrap Exploris 480 (12 SPD) or Orbitrap Astral (16 and 50 SPD). The corresponding protein ion intensities and unique peptide counts are provided in Tables S1-1 and S1-2, respectively. (B–D) Numbers of detected proteins associated with OMIM (B), RUSP (C), and recessive disease genes included in BEACONS (D) under each condition. In panels A–D, data are shown as the mean ± SD of the corresponding protein counts for individual samples (*n* = 4 per condition). (E) Reproducibility of DBS analysis using the EDTA-enhanced NANDA workflow, assessed using 16 independently prepared DBS samples subjected to the 100 mM EDTA wash and analyzed using an Orbitrap Astral at 50 SPD under identical LC–MS settings. Pairwise Pearson correlation coefficients were calculated from protein ion intensities obtained with MBR enabled. Abbreviations: DBS, dried blood spot; EDTA, ethylenediaminetetraacetic acid; NANDA, non-targeted analysis of non-specifically DBS-absorbed proteins; SD, standard deviation; SPD, samples per day; OMIM, Online Mendelian Inheritance in Man; RUSP, Recommended Uniform Screening Panel; BEACONS, Building Evidence and Collaboration for GenOmics in Nationwide Newborn Screening; LC–MS, liquid chromatography–mass spectrometry.

The 50 SPD method was employed for high-throughput analysis. At 50 SPD and 70% instrument uptime, the theoretical annual capacity is approximately 12,800 measurements. A conservative estimate of approximately 10,000 study samples per year allows additional capacity for quality-control measurements, repeat analyses, and scheduling. Using the Astral 50 SPD method, 6,187 proteins were identified while reducing the analysis time to approximately 27% of that required for the Exploris 12 SPD method (Figure 7A). This combined sample-preparation and acquisition workflow yielded more identifications than conventional NANDA analyzed with Exploris 12 SPD; the specific effect of EDTA was assessed separately in the matched Exploris comparison. Among the identified proteins, the numbers of OMIM-related proteins were 2,165 with Exploris 12 SPD, 2,400 with Astral 16 SPD, and 2,133 with Astral 50 SPD (Figure 7B). In addition, the numbers of Recommended Uniform Screening Panel-related proteins^28^ were 45, 51, and 46, respectively (Figure 7C), and the Building Evidence and Collaboration for GenOmics in Nationwide Newborn Screening-related proteins encoded by a recessive allele (https://www.beaconsnbs.org/) were 275, 297, and 276, respectively (Figure 7D). Under matched Exploris 12 SPD conditions, EDTA-enhanced sample preparation increased OMIM-associated protein identifications from 1,934 to 2,165, RUSP-associated identifications from 42 to 45, and BEACONS-associated identifications from 251 to 275 (Figure 7B–D; Table S1). Disease-associated coverage was also largely retained with the high-throughput Astral 50 SPD method. In a cross-platform serum proteomics study, Katsavelis et al. demonstrated that sample preparation shaped the detected protein repertoire and associated functional profiles even when advances in MS instrumentation improved analytical depth.^33^ Although serum and DBS differ in sample composition and processing requirements, this finding highlights the importance of sample preparation when interpreting proteome coverage. The increased detection of OMIM-, RUSP-, and BEACONS-associated proteins under matched Exploris 480 conditions demonstrates that EDTA-enhanced sample preparation expands disease-associated protein coverage independently of an instrument upgrade. Finally, reproducibility was evaluated using 16 independently prepared DBS samples processed using the EDTA-enhanced NANDA workflow and analyzed using the Astral 50 SPD method. Both the mean and median Pearson correlation coefficients for the LC–MS/MS data were 0.996 (Figure 7E). This demonstrated high reproducibility across the independently prepared samples evaluated here.

### Mouse DBS Proteomics Reveals Shared and Disease-Specific Immune Signatures in Inflammatory Mouse Models

Considering that DBS can be prepared from a small volume of blood, the EDTA-enhanced NANDA workflow was applied to small experimental animals (mice). To characterize the systemic immune signatures detectable in peripheral blood, proteomic profiling of DBS prepared from the whole blood of control mice and two inflammatory disease models (EAE and AD) was performed.

The depletion of high-abundance soluble proteins using the EDTA-enhanced NANDA workflow prior to analysis enabled the identification of >7,000 proteins with Astral 16 SPD, including proteins with lower signal intensities (Figure 8A and Table S2-1). Mean identification counts were 7,849 in control, 7,094 in EAE, and 7,380 in AD mice (*n* = 4 per group). Residual fibrinogen-chain and hemoglobin ion intensities were also compared across the three groups (Figure S3). Fibrinogen-chain signals did not differ significantly between either disease group and control, whereas hemoglobin signal was higher in EAE than control (*p* < 0.01) and did not differ significantly between AD and control. The higher residual hemoglobin signal in EAE could contribute to the lower protein identification count, so a contribution from differences in residual abundant proteins cannot be excluded. The resulting proteomic profiles clearly differed between the disease models (Figure 8B). Compared with proteins in control mice, 451 and 298 proteins were upregulated and downregulated in EAE-induced mice, respectively, whereas 159 and 594 proteins were upregulated and downregulated in AD-induced mice, respectively (Figure 8C, D). Although a subset of proteins showed shared alterations between the two disease models, the overlap depended on the direction of change (Figure 8E, F). Among increased proteins, 124 were shared, 327 were specific to EAE, and 35 were specific to AD. Among decreased proteins, 203 were shared, 95 were specific to EAE, and 391 were specific to AD. Because starting abundances and depletion efficiencies were not independently measured, these data do not establish a depletion failure or a purely biological explanation for the group differences.

**Figure 8.**
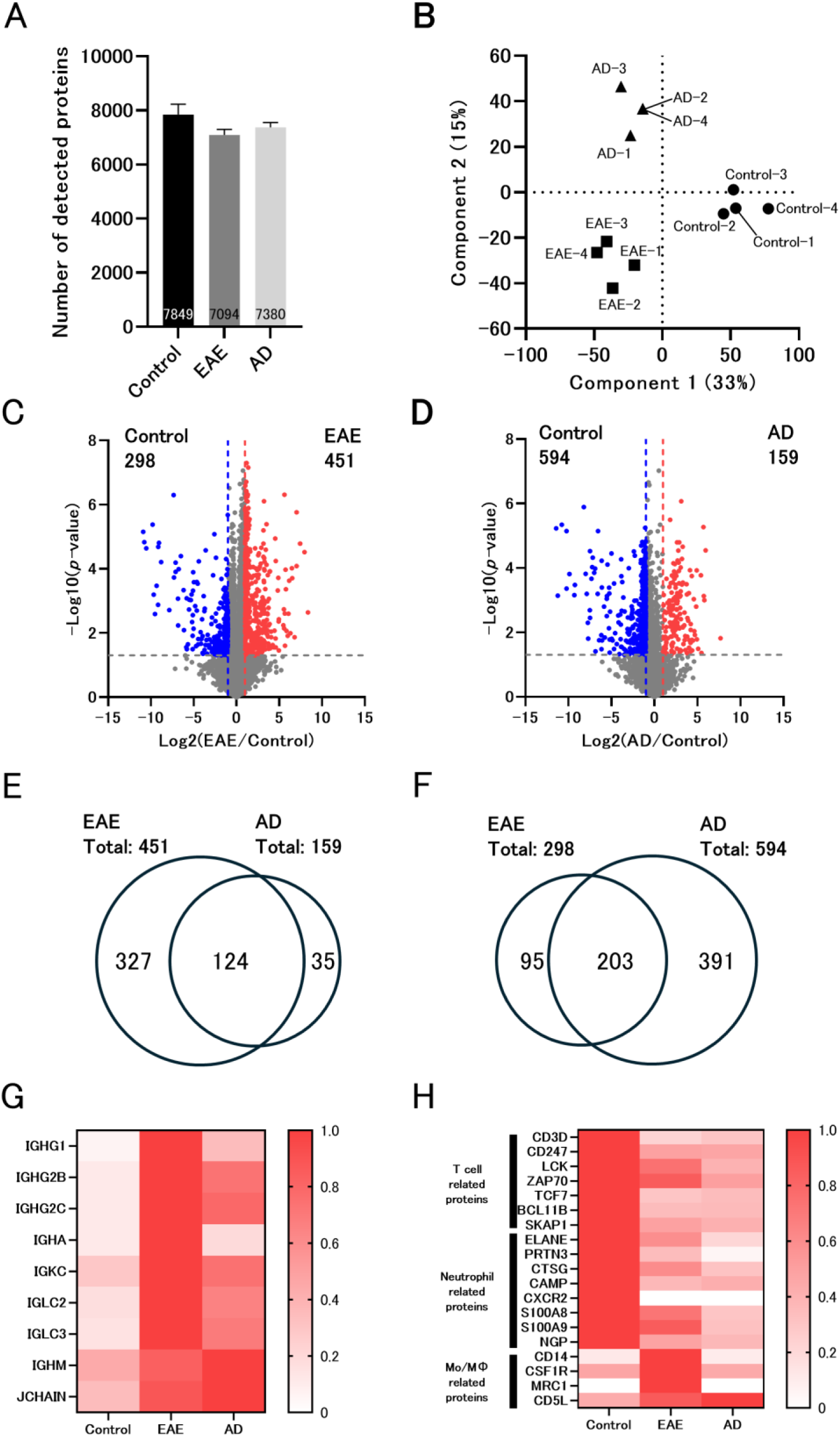
Comparative proteomic profiling of control, EAE, and AD model mice using an EDTA-enhanced NANDA workflow. (A) Number of proteins detected in DBS from control, EAE, and AD mice, shown as the mean ± SD of protein counts for individual samples (*n* = 4 per group). Counts were obtained from independent analysis of individual runs (Table S2-1). DIA LC–MS/MS analysis was performed using an Orbitrap Astral at 16 SPD. Panels B–H use the jointly analyzed dataset with cross-run normalization and without MBR (Table S2-2). (B) Principal component analysis (PCA) of detected proteins showing separation among the control, EAE, and AD groups. Circles, squares, and triangles represent control, EAE, and AD mice, respectively. Individual sample labels identify all four mice in each group, including overlapping AD samples. (C–D) Volcano plots comparing protein abundances in EAE (C) and AD (D) mice with those in control mice (*n* = 4 per group). Proteins with |log₂(fold change)| ≥ 1 and a two-tailed, unpaired Welch’s *t*-test *p*-value < 0.05 were considered significantly altered. Red and blue dots indicate proteins with significantly higher abundance in the disease and control groups, respectively. The x-axis represents log₂(disease/control fold change), and the y-axis represents −log₁₀(*p*-value). (E–F) Venn diagrams comparing proteins significantly increased (E) or decreased (F) in EAE and AD mice relative to control mice, using the criteria specified for panels C and D. (G–H) Heatmaps showing relative abundances of immune-related proteins across the control, EAE, and AD groups. For each protein, the mean ion intensity (*n* = 4 per group) was normalized to the highest group mean. (G) Humoral immunity-associated proteins, including immunoglobulin heavy and light chains and JCHAIN, showing increased abundance in both disease models. (H) Immune signatures distinguishing the models, including increased monocyte/macrophage-associated proteins in EAE and reduced signals for T-cell- and neutrophil-associated proteins, with patterns differing between models. Protein names are presented as labeled in the figure; all samples were obtained from mice. Abbreviations: DBS, dried blood spot; EAE, experimental autoimmune encephalomyelitis; AD, atopic dermatitis; EDTA, ethylenediaminetetraacetic acid; NANDA, non-targeted analysis of non-specifically DBS-absorbed proteins; DIA, data-independent acquisition; LC–MS/MS, liquid chromatography–tandem mass spectrometry; PCA, principal component analysis; SD, standard deviation; SPD, samples per day.

A common feature observed in both EAE and AD-induced mice was the increased abundance of proteins associated with humoral immunity, including immunoglobulins (Figure 8G and Table S2-2). Notably, the composition of the immunoglobulin-related proteins differed between the two models. Class-switched immunoglobulin heavy and light chain components, including IGHG1, IGHG2B, IGHG2C, IGHA, IGKC, IGLC2, and IGLC3, were more abundant in the EAE group than in the AD group. In contrast, IGHM and JCHAIN levels increased in AD-induced mice. These differences suggest model-dependent variation in immunoglobulin-chain composition, with relatively higher levels of several class-switched immunoglobulin components in EAE and increased IGHM- and JCHAIN-associated signals in AD.

Given that JCHAIN is required for the formation of pentameric IGM and dimeric IGA, increased levels of IGHM and JCHAIN in AD may reflect ongoing or barrier-associated humoral immune responses. In contrast, several proteins involved in T-cell receptor signaling were decreased in both EAE and AD, including CD3D, CD247, LCK, ZAP70, TCF7, BCL11B, and SKAP1. This indicates a reduced T-cell-associated signature in the peripheral blood. Proteins associated with neutrophil biology, including ELANE, PRTN3, CTSG, CAMP, CXCR2, S100A8, S100A9, and NGP, were also markedly downregulated (Figure 8H and Table S2-2).

Considering that these proteins represent well-established neutrophil granule components and inflammatory markers, their coordinated reduction suggests a diminished neutrophil-associated signature in peripheral blood. Previous studies have implicated neutrophils in inflammatory processes associated with multiple sclerosis and its experimental models, as well as atopic dermatitis.^34,35^ However, the reduced abundance of T-cell- and neutrophil-associated proteins in DBS does not directly demonstrate migration of these cells from the circulation into affected tissues. Such changes may also reflect differences in circulating blood-cell composition or protein abundance within individual cell populations. Therefore, additional analyses, including blood-cell phenotyping and tissue-level examinations, would be required to determine the mechanisms underlying these observations.

In the EAE model, several proteins associated with monocyte and macrophage lineages were increased, including CD14, CSF1R, and MRC1, which suggests the activation of monocyte/macrophage-related immune pathways (Figure 8H). CD5L abundance was increased in both EAE and AD mice relative to control mice, indicating that this change was shared between the two models (Figure 8H). The adhesion molecule ICAM1 also increased, consistent with an altered leukocyte adhesion-associated signature (Table S2-2). In addition, immunoproteasome subunit (PSMB8 and PSMB10) levels were increased, which is consistent with the activation of interferon-responsive antigen-processing pathways during autoimmune inflammation. In the AD proteomic profile, epithelial or secretory-associated proteins, such as SCGB1A1 and keratin family members, including KRT73, were detected. This possibly reflects the release of tissue-derived proteins during inflammatory skin processes (Table S2-2). These findings can be compared with previous studies of the corresponding disease models. In MOG35-55-induced EAE, antigen-specific IgG1 responses have been reported following immunization.^36^ Previous proteomic analysis of peripheral blood mononuclear cells from EAE mice also revealed extensive disease-associated changes in immune-related proteins.^37^ Furthermore, CSF1R-dependent myeloid cells have been implicated in EAE pathogenesis,^38^ providing biological context for the increased abundance of CSF1R and other myeloid-associated proteins observed in our study. In MC903-induced dermatitis, previous studies demonstrated increased serum IgE and IgG levels and type 2 inflammation,^19^ as well as neutrophil infiltration into inflamed skin.^39^ These findings are broadly consistent with the humoral and immune-cell-associated signatures detected in our DBS samples. However, direct comparisons of individual protein changes are limited by differences in sample types, analytical methods, and disease conditions among studies. Moreover, changes in immunoglobulin-chain abundance or immune-cell-associated proteins in whole-blood DBS do not directly establish antibody specificity, B-cell maturation states, or immune-cell redistribution.

Collectively, these findings suggest that EAE is characterized by relatively higher levels of several class-switched immunoglobulin components and increased monocyte/macrophage-associated proteins, whereas AD exhibits increased IGHM- and JCHAIN-associated signals and more pronounced reductions in T-cell- and neutrophil-associated proteins in peripheral blood. These results demonstrate the ability of DBS proteomics to detect shared and distinct immune-associated proteomic patterns in inflammatory mouse models.

## CONCLUSION

This study developed an EDTA-enhanced NANDA workflow that enables deep and high-throughput proteomic profiling of DBS. The incorporation of an EDTA wash reduced residual fibrinogen-chain signals and improved proteome coverage; the hemoglobin response depended on EDTA concentration, and 100 mM was selected for overall coverage and compatibility with standard DBS collection rather than maximal hemoglobin removal. Under matched Exploris 12 SPD conditions, the improved sample preparation also expanded detection of disease-associated proteins, demonstrating a contribution independent of the instrument upgrade. Although EDTA chelates divalent metal ions, the precise molecular mechanism underlying the enhanced removal of abundant proteins remains to be determined. Importantly, this improvement was achieved without requiring any modification of the standard DBS collection procedures, thereby ensuring compatibility with existing clinical workflows. When combined with the Orbitrap Astral DIA-MS, the optimized workflow enabled the identification of >7,000 proteins in a single-shot analysis. To the best of our knowledge, this represents the deepest proteome coverage reported for DBS to date. In addition to enhanced depth, the workflow demonstrated high reproducibility and analytical robustness. These features support its applicability to high-throughput workflows, including settings relevant to NBS, where both scalability and consistency are critical. Furthermore, the application of this approach to mouse disease models demonstrated that DBS proteomics can capture systemic immune alterations from minimal blood volumes. The ability to detect both shared and disease-specific immune signatures highlights the potential of this platform for achieving minimally invasive longitudinal monitoring of disease progression and immune dynamics in small-animal studies. Overall, the EDTA-enhanced NANDA workflow overcomes key limitations of DBS proteomics by improving proteome depth while maintaining simplicity, scalability, and clinical compatibility. Therefore, this platform provides a practical and broadly applicable solution for deep proteomic analysis in clinical, epidemiological, and experimental settings.

## ASSOCIATED CONTENT

### Supporting Information

GO enrichment analyses for molecular function, biological process, and cellular component of proteins detected exclusively with the 100 mM EDTA wash; comparison of protein coverage and ion-intensity correlations between the 100 mM EDTA wash and 500 mM EDTA-impregnated filter paper; and fibrinogen and hemoglobin ion intensities in control, EAE, and AD mouse DBS (Figures S1–S3) (PDF).

Protein ion intensities and unique peptide counts in human DBS across NANDA workflows and LC–MS conditions, with OMIM, RUSP, and BEACONS annotations (Table S1) (XLSX).

Protein groups and ion intensities in control, EAE, and AD mouse DBS from independent analysis of individual runs and joint analysis of all 12 runs (Table S2) (XLSX).

### Availability of data and materials

The LC–MS/MS data supporting the conclusions of this article are available from the ProteomeXchange Consortium in the jPOST partner repository, under the accession code PXD081089

## AUTHOR INFORMATION

### Author Contributions

D.N. and Y.K. designed the study; D.N. performed the research; T.K. and N.U. provided reagents and samples; H.M., Y.O. and R.K. analyzed the data; D.N., Y.E., O.O., and Y.K. wrote the paper with input from all authors.

## ABBREVIATIONS

ACN: acetonitrile
AD: atopic dermatitis
AGC: automatic gain control
ATP: adenosine triphosphate
auto-NANDA: automated NANDA
BEACONS: Building Evidence and Collaboration for GenOmics in Nationwide Newborn Screening
BP: biological process
CC: cellular component
DAVID: Database for Annotation Visualization and Integrated Discovery
DBS: dried blood spots
DIA: data-independent acquisition
DIA-MS: data-independent acquisition mass spectrometry
DMNG: decyl maltose neopentyl glycol
EAE: experimental autoimmune encephalomyelitis
EDTA: ethylenediaminetetraacetic acid
EtOH: ethanol
GO: Gene Ontology
GPI: glycosylphosphatidylinositol
GTP: guanosine triphosphate
HGNC: HUGO Gene Nomenclature Committee
IM: ion mobility
LC: liquid chromatography
LC–MS: liquid chromatography–mass spectrometry
LC–MS/MS: liquid chromatography–tandem mass spectrometry
LMNG: laurylmaltose neopentylglycol
MBR: match-between-runs
MC903: calcipotriol
MF: molecular function
Mo/MΦ: monocytes/macrophages
MS: mass spectrometry
MW: molecular weight
NANDA: non-targeted analysis of non-specifically DBS-absorbed proteins
NBS: newborn screening
NNs: neural networks
ns: not significant
OMIM: Online Mendelian Inheritance in Man
PCA: principal component analysis
RNA: ribonucleic acid
RT: retention time
RUSP: Recommended Uniform Screening Panel
SD: standard deviation
SDB: styrene–divinylbenzene
SH2: Src homology 2
SPD: samples per day
TBST: Tris-buffered saline with Tween 20
TFA: trifluoroacetic acid
UHPLC: ultra-high-performance liquid chromatography
VAMS: volumetric absorptive microsampling.

