## Supplementary material for "Development of an Ethylenediaminetetraacetic Acid-Enhanced Deep Proteomic Profiling Method for Dried Blood Spots and its Application in Mouse Disease Models": Figure S1

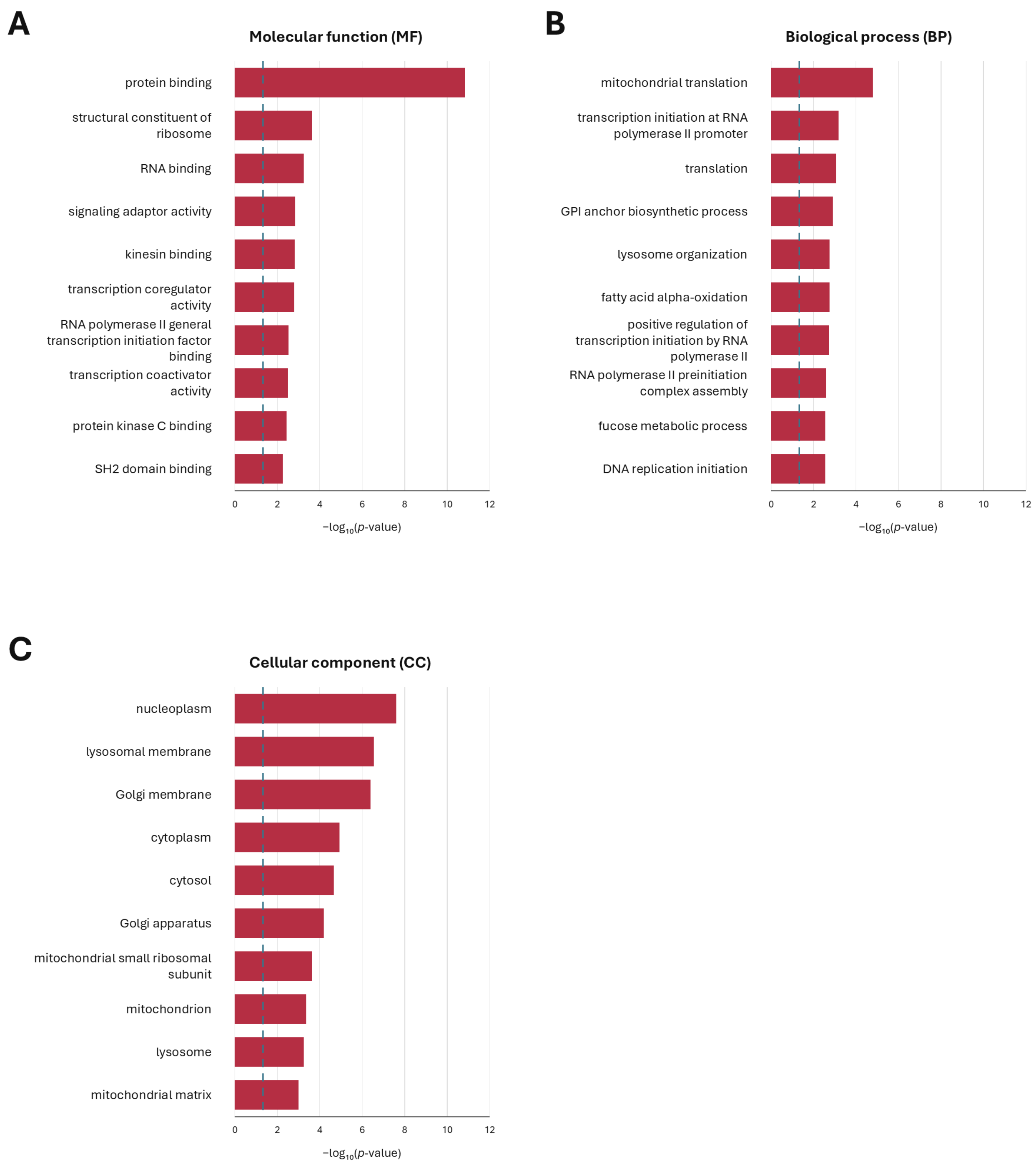


Figure S1. GO enrichment analyses of the 773 proteins detected exclusively in the 100 mM EDTA wash condition. (A) Molecular function (MF). (B) Biological process (BP). (C) Cellular component (CC). The top 10 terms in each category, ranked by *p*-value, are shown. Bars represent $-\log_{10}(p\text{-value})$; the dashed line indicates $p = 0.05$.
