## Supplementary material for "Development of an Ethylenediaminetetraacetic Acid-Enhanced Deep Proteomic Profiling Method for Dried Blood Spots and its Application in Mouse Disease Models": Figure S2

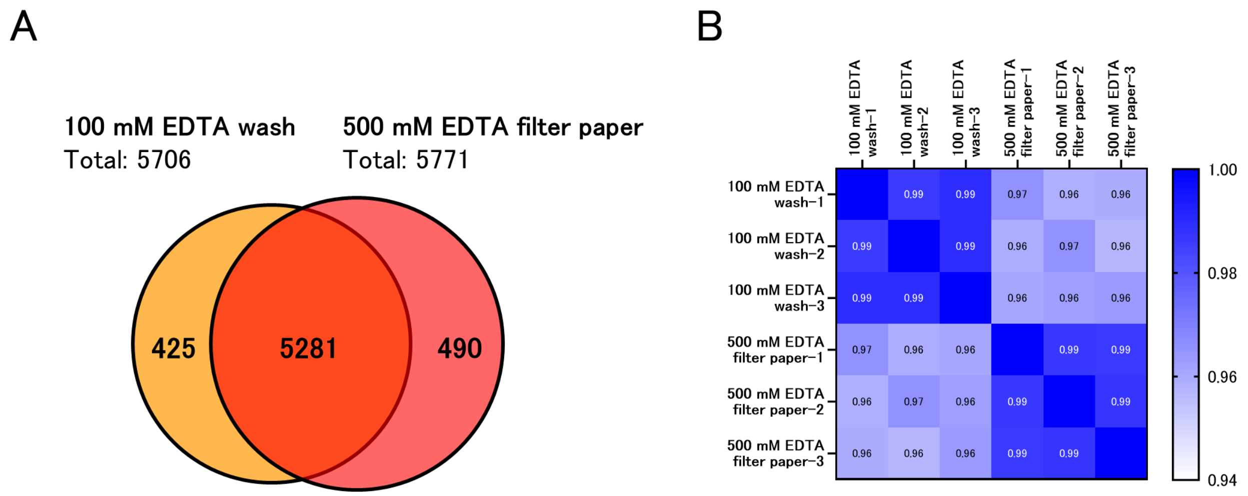


Figure S2. Comparison of DBS proteomic profiles obtained using a 100 mM EDTA wash or filter paper impregnated with 500 mM EDTA. (A) Venn diagram comparing proteins detected in all three replicates within each condition (*n* = 3 per condition). Of these, 5,281 proteins were shared between conditions, 425 were detected only under the 100 mM EDTA wash condition, and 490 were detected only under the 500 mM EDTA-impregnated filter paper condition. (B) Heatmap of pairwise correlation coefficients of protein ion intensities across individual samples (*n* = 3 per condition), using data analyzed with MBR enabled. DIA LC–MS/MS analysis was performed using an Orbitrap Exploris 480 at 12 SPD. Abbreviations: DBS, dried blood spot; EDTA, ethylenediaminetetraacetic acid; SPD, samples per day.
