## Supplementary material for "Development of an Ethylenediaminetetraacetic Acid-Enhanced Deep Proteomic Profiling Method for Dried Blood Spots and its Application in Mouse Disease Models": Figure S3

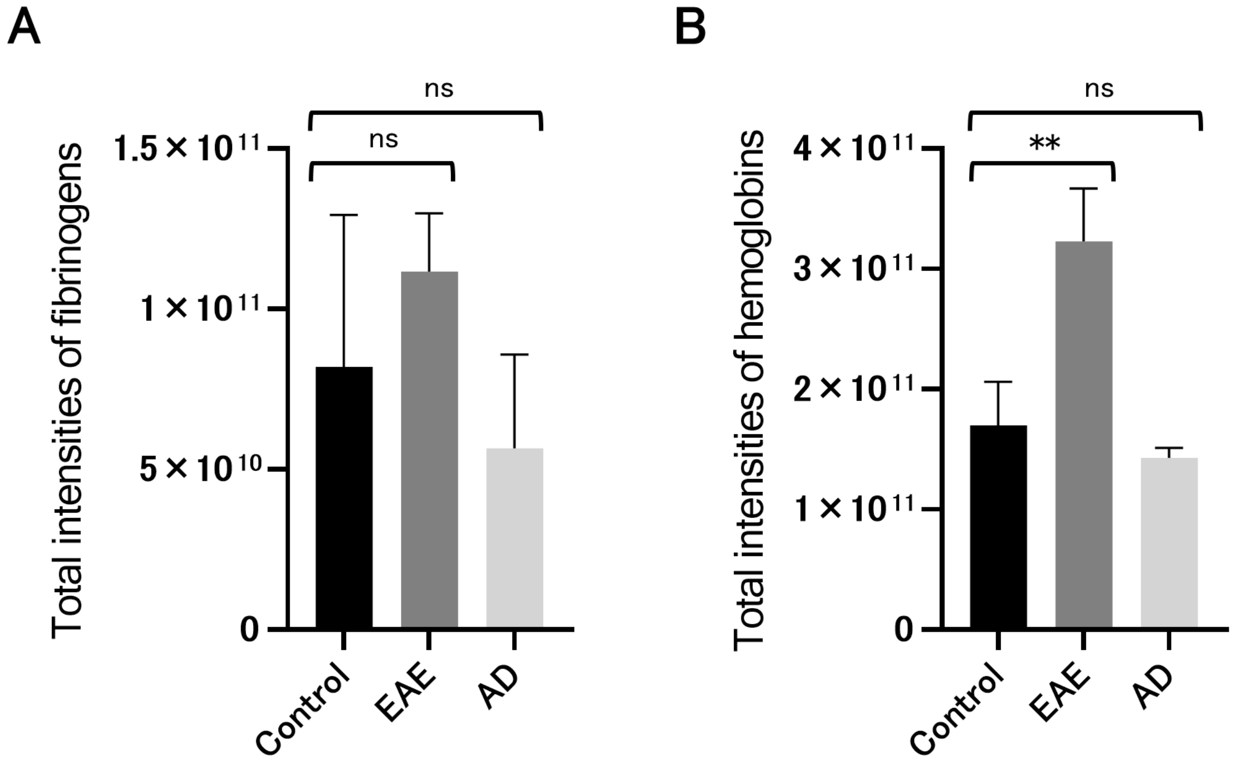


Figure S3. Comparison of fibrinogen and hemoglobin ion intensities in DBS samples from control, experimental autoimmune encephalomyelitis (EAE), and atopic dermatitis (AD) mice processed using EDTA-enhanced NANDA. (A) Summed ion intensities of fibrinogen chains (Fga, Fgb, and Fgg). (B) Summed ion intensities of hemoglobin subunits (Hba, Hbb-b1, Hbb-b2, Hbb-bh1, Hbb-bs, Hbb-y, Hbz, and Hbq1b). Data are shown as the mean ± SD ($n$ = 4 mice per group). Statistical significance was assessed against the control using two-tailed, unpaired Welch's $t$-tests: **$p < 0.01$; ns, not significant. DIA LC–MS/MS analysis was performed using an Orbitrap Astral at 16 SPD. Abbreviations: DBS, dried blood spot; EDTA, ethylenediaminetetraacetic acid; NANDA, non-targeted analysis of non-specifically DBS-absorbed proteins; SD, standard deviation; SPD, samples per day.
